# Bayesian phylodynamics of avian influenza virus H9N2 in Asia with time-dependent predictors of migration

**DOI:** 10.1101/450064

**Authors:** Jing Yang, Nicola F. Müller, Remco Bouckaert, Bing Xu, Alexei J. Drummond

**Author notes:** Corresponding authors (JY); (BX); (AJD).

## Abstract

Model-based phylodynamic approaches recently employed generalized linear models (GLMs) to uncover potential predictors of viral spread. Very recently some of these models have allowed both the predictors and their coefficients to be time-dependent. However, these studies mainly focused on predictors that are assumed to be constant through time. Here we inferred the phylodynamics of H9N2 viruses isolated in 12 Asian countries and regions under both discrete trait analysis (DTA) and structured coalescent (MASCOT) approaches. Using MASCOT we applied a new time-dependent GLM to uncover the underlying factors behind H9N2 spread. We curated a rich set of time-series predictors including annual international live poultry trade and national poultry production figures. This time-dependent phylodynamic prediction model was compared to commonly employed time-independent alternatives. Additionally the time-dependent MASCOT model allowed for the estimation of viral effective sub-population sizes and their changes through time and these effective population dynamics within each country were predicted by a GLM. International annual poultry trade is a strongly supported predictor of virus migration rates. There was also strong support for geographic proximity as a predictor of migration rate in all GLMs investigated. In time-dependent MASCOT models, national poultry production was also identified as a predictor of virus genetic diversity through time and this signal was obvious in mainland China and Bangladesh. Our application of a recently introduced time-dependent GLM predictors integrated rich time-series data in Bayesian phylodynamic prediction. We demonstrated the contribution of poultry trade and geographic proximity (potentially unheralded wild bird movements) to avian influenza spread in Asia. To gain a better understanding of the drivers of H9N2 spread, we suggest increased surveillance of the H9N2 virus in countries that are currently under-sampled as well as in wild bird populations in the most affected countries.

**Author summary:** What drives the geographic dispersal and genetic diversity of H9N2 avian influenza virus in Asia? We used two model-based approaches, DTA and MASCOT, to reconstruct the phylogeographic dynamics of the virus. Further, multiple potential predictors were used to inform the virus spread and population dynamics by GLMs. Here, we maximised the power of time-series predictors in Bayesian phylodynamic prediction. For the first time, we were able to quantify the contribution of both time-series and constant predictors to both migration rates and effective population sizes in a structured population. We identified a positive association of international poultry trade and national poultry production time-series with virus migration rates and effective population sizes respectively. We also identify geographic proximity as a strongly supported driver to virus migration rates and this points to the potential role of wild bird populations in virus dispersal across countries. Our study is a practical exemplar of the use of temporal information in predictors to model heterogeneous spatial diffusion and population dynamic processes and provides direction to H9N2 control efforts in Asia.

## Introduction

### Phylogeographic models and their extended GLMs

To infer the migration history of sampled lineages based on genetic data, phylogeographic methods can be used. The discrete trait analysis (DTA) and structured coalescent model are commonly used probabilistic model-based phylogeographic methods. The DTA model treats the migration of lineages between different geographic locations as a per-lineage continuous-time Markov process, analogous to the DNA substitution process (Lemey et al., 2009). This approach achieves computational efficiency by integrating over all possible migration histories using the efficient tree pruning algorithm for computing phylogenetic likelihoods (Felsenstein, 1981). One drawback of this approach is the assumption of independence of the phylogenetic tree generating process from the migration process, which can lead to bias and underuse of the data (De Maio et al., 2015). Another drawback is the sensitivity of such models to local sampling intensity, since such a model assumes that the sample sizes across subpopulations are proportional to the sub-population sizes. As a result biases are expected in the estimated migration rates for data sets where sampling is biased across sub-populations (De Maio et al., 2015). An alternative approach that is robust to variation in sample sizes is the structured coalescent, which describes the genealogy of individuals randomly sampled from a structured population and thereby coherently models the interaction between migration and coalescent processes (Takahata, 1988). In such coalescent methods the main assumption is that the sample size is small compared to the population size. It is further assumed that individual subpopulations are well mixed. Exact inference under the structured coalescent is challenging (Vaughan et al., 2014). Thus, approximations to the structured coalescent model by approximately integrating over all ancestral migration histories were proposed (De Maio et al., 2015; Müller et al., 2018). The marginal approximation of the structured coalescent (MASCOT) currently provides the closest approximation to the structured coalescent while being computationally efficient. This allows to analyse datasets with many different sub-populations (Müller et al., 2017, 2018).

In order to inform phylogeographic reconstructions from predictor data, generalized linear models (GLMs) can be employed to inform the pathogen migration rates between distinct geographical locations (Lemey et al., 2014). Some authors used GLMs in both discrete and continuous phylogeographic models to investigate the impact of underlying environmental variables on the dispersal frequencies and velocity of a virus respectively (Dellicour et al., 2018). But only univariate models can be considered in the current GLM implementation in continuous phylogeographic inference. The GLM model incorporating multiple predictors in DTA (Lemey et al., 2014) gained popularity for its computational efficiency and user-friendly implementation in BEAST 1.10 (Suchard et al., 2018). But it unrealistically assumes time-homogeneous substitution processes between sub-populations. To allow for time-dependence in both coefficients and predictors data, the epoch GLM has already been proposed to model the heterogeneous spatial diffusion processes through time in DTA (Dudas et al., 2017; Bielejec et al., 2014). However, only time-dependent coefficients (rather than time-dependent predictors) were considered in a recent phylodynamic GLM to inform the temporal dynamics in the spread of Ebola virus (Dudas et al., 2017). Very recently the GLM model with both time-dependent predictors and coefficients were proposed in MASCOT (Müller et al., 2018). This allows, for the first time, the ability to quantify the contribution of both time-series and constant predictors to both migration rates and effective population sizes jointly in a structured population.

### Avian influenza H9N2 in Asia

H9N2 avian influenza viruses (AIVs) have spread into multiple Asian countries and became endemic in domestic poultry populations in some of these countries (Guan et al., 1999; Cameron et al., 2000; Nili and Asasi, 2003). Its transnational geographic dispersal and exchange of genetic segments with other subtypes in poultry increase their potential for zoonotic threat to public health (Guan et al., 1999; Liu et al., 2013; Qi et al., 2014). Of note, the multi-segmented H9N2 virus provided its internal gene materials to facilitate the genesis of the novel H7N9 AIVs that caused multiple outbreaks and high mortality in humans since 2013 in China (Liu et al., 2013). Hereafter the underlying mechanism behind evolution and spread of the ”donator” H9N2 virus raises wide concerns (Pu et al., 2015).

The poultry trade network is a potential source of avian influenza virus mobility in Asia and may help to explain the multiple introductions of H9N2 AIVs from the same genetic group into different countries (Bahl et al., 2016; Hu et al., 2017). The G1-like H9N2 virus isolated in middle eastern countries shares a common ancestor with the virus from China (Shanmuganatham et al., 2016) despite the lack of an east-west migratory flyway in Asia. Additionally, genetically related H9N2 viruses can be found in geographic regions separated by long distances, which suggests a role for migration by poultry trade. Further, the asymptomatic poultry carrying this low pathogenetic virus could be neglected during transportation process. Poultry transportation can bring together various host species from different regions in a high-density setting and provides an ideal environment for interspecies virus transmission and theretofore the reassortment of different viral segments (Choi et al., 2005).

Another potential source of virus mobility is wild bird movements. Wild aquatic birds can act as natural reservoirs of influenza viruses, since infection is often asymptomatic, allowing them to migrate while carrying the virus (Keawcharoen et al., 2008). Viral transmission from domestic to wild birds will occur within a region while mobile and migratory wild birds could transmit viruses between regions (Bahl et al., 2016). Although poultry trade may play a major role in H9N2 virus spread, the contribution of wild bird populations to the long distance virus dispersal to other poultry populations should therefore not be neglected. Nevertheless, we here focus on assessing the role of poultry trade and production in the prevalence and spread of H9N2 AIVs.

By elucidating the evolutionary dynamics and the underlying factors that drive virus spread we aim to better understand the ecology and spread mechanism of the viruses in Asia. In this study, the spatial distribution and temporal dynamic of H9N2 viruses were described to understand their basic conditions. The migration dynamics and their underlying mechanism of H9N2 AIVs between 12 Asian countries and regions were inferred by using two phylogeographic methods DTA and MASCOT in a Bayesian Markov chain Monte Carlo (MCMC) inference framework. Under MASCOT, we also jointly inferred predictors of the effective population size of the virus in each location, which is not possible in DTA. To do so, we used a GLM approach to parameterize migration rates and effective population sizes of H9N2 viruses by potential predictors, including time-dependent predictors (annual live poultry trade, annual national poultry production, yearly mean temperature, yearly total rainfall, annual seasonality of temperature and rainfall, and yearly virus sample size) and time-independent predictors (e.g. geographic distance). The underlying mechanisms of virus migration and genetic diversity were evaluated and the sensitivity of our results was investigated by comparing models that included different predictors, different priors on the inclusion probability of predicator and alternative phylogeographic modelling assumptions.

## Materials and methods

### Sequence data

We obtained all full-length haemagglutinin (HA) segment nucleotide sequences of avian-origin H9N2 from Asian countries and regions that were available in the GenBank Influenza Virus Database (http://www.ncbi.nlm.nih.gov/genomes/FLU/FLU.html) hosted by the National Center for Biotechnology Information (NCBI). Identical sequences caused by duplicated submissions in the database (i.e. same sequence and same isolate name), were reduced to a single sequence to avoid bias in rate estimates. Sequences without explicit isolation date or country information were excluded. These HA sequences from 1976 to 2014 were geocoded and pooled into groups according to their geographic location, host type (chicken, duck, quail, turkey, wild bird, and others), and isolation date (S4 Table).

The virus sequences were aligned in MAFFT v7 software with default parameters (Katoh and Standley, 2013). Recombination was analyzed in RDP v4.6.3 software (Martin et al., 2015) and recombinant sequences identified by more than half of selected recombination detection methods were removed. We evaluated the temporal signal of the remaining heterochronous sequences with TempEst (Rambaut et al., 2016) and removed sequences that we identified as outliers. To get a more even distribution of samples through time and between different locations, we randomly sub-sampled the H9N2 sequences to keep at most 10 isolates per country per year. In order to avoid overparameterization we discarded locations with less than 10 isolates in total. Finally, we added commonly used representative HA gene sequences to help the phylogenetic clade classification (Guo et al., 2000). The final data set contained 526 HA sequences from 12 Asian countries/locations. We defined distinct locations as Bangladesh, mainland China, Hong Kong, South Korea, Japan, Vietnam, Indian, Pakistan, Iran, Israel, Saudi Arab, and the United Arab Emirates. More detail information on the viral sequences is provided in S4 Table.

### Empirical predictors

To inform the spread and genetic diversity of H9N2 viruses across the 12 locations, we chose several potential predictors. These are similar to ones previously used to describe the spread of H3N2 (Lemey et al., 2014) or Ebola (Dudas et al., 2017; Müller et al., 2018). We used annual live poultry trade, annual poultry production, gross domestic product values (GDP), geographic distance, a predictor describing if countries share a continental border, temperature, temperature seasonality, rainfall, rainfall seasonality, virus sample size and the latitude of centroid point of each country. All the country-level predictors were available from 1986 to 2013.

The live poultry trade and poultry production data (including data related to chicken, duck, and turkey) were downloaded from FAOSTAT (http://faostat3.fao.org). Typically, poultry movements are driven by variation in the supply and demand for poultry, which are in turn commonly affected by economic, ecological and climatic conditions (Nicolas et al., 2018). These, as well as practical considerations led us to chose the set of potential predictors used in this study.

GDP statistics were collected to describe the economic level of each country. Annual mean temperature and annual total rainfall were gathered to describe the climatic condition in each country through time. Further, the annual variation of temperature and rainfall were described by temperature seasonality and rainfall seasonality respectively. Temperature seasonality is the standard deviation of the monthly mean temperatures in each year. Rainfall seasonality (*R_S_*) for year *t* is the ratio of the standard deviation of the monthly total precipitation (*s_p_*) over one plus the mean monthly precipitation (*p_m_*): *R_S_*(*t*) = *s_p_*(*t*)/(1 + *p_m_*(*t*)). GDP, temperature, and rainfall data were collected from the World Bank database (http://data.worldbank.org/).

The H9N2 isolates sampled from the same region commonly tended to gather in a phylogenetic group. The geographic distance between each pair of locations was considered a potential factor of virus spread and calculated by the great circle distance based on the central latitude and longitude of each location. We also used a predictor with 1 or 0 to describe if two locations share their border on the continent or not, respectively. To test the impact of sampling effects, the number of H9N2 sequences in both origin and destination location was considered as two separate predictors. Finally the geographic central latitudes of locations were considered as a predictor to investigate latitude as a potential driver of H9N2 spread.

To avoid excessive co-linearity among explanatory predictors, we removed the GDP, temperature seasonality and the latitude variables, since the Pearson correlation coefficients between temperature variable and each of them exceeded 0.7. To eliminate the effect of the magnitude of different predictors, all predictors (except binary predictors) were transformed into log space and standardized so that their means are equal to 0 and standard deviation equals to 1. This is standard practice when using generalize linear models to inform viral migration rates and effective population size (e.g. (Lemey et al., 2014)). More detail information on predictors is supplied in S1 Table.

## Methods

### The DTA model

The DTA model treats movement of viral genes across discrete geographic locations as a continuous time Markov process in which the state space consists of the sampled locations (Lemey et al., 2009; Drummond and Bouckaert, 2015). This model treats the spread of viruses as statistically equivalent to the evolution of molecular substitutions at a site. The posterior probability distribution of parameters given data in the DTA model is shown in Eq 1 (Lemey et al., 2009). Here, the aligned sequences *S* and the sampling locations *L* are treated as observations, whereas the isolation dates of the sequence *t* are treated as boundary conditions. The phylogenetic tree *T*, the nucleotide substitution rate matrix *µ*, the forwards-in-time migration rate matrix f and the effective population size *θ* of the whole meta-population are random variables estimated in this model. The first term on the right is the likelihood of the sequences. This likelihood is calculated by integrating over all possible substitution histories using the pruning algorithm (Felsenstein, 1981). The second term is the likelihood of the sampling locations given the time-stamped genealogy and the instantaneous migration rate matrix. It is calculated by the same pruning algorithm, but using a migration rate matrix rather than a substitution matrix. The third term describes the probability density of the genealogy across the entire meta-population, approximated by a standard neutral coalescent prior for a well-mixed and unstructured population. The fourth term represents the prior distribution of three independent random variables. It should be noted that to the extent that *θ* can be interpreted in this model, it represents the effective population size for the entire meta-population across all locations in the dataset.

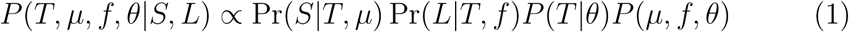

### The structured coalescent model

The structured coalescent jointly models how lineages coalesce within locations and migration between them. The posterior distribution of the parameters given the data in a structured coalescent phylogeographic inference is described in Eq 2. Here, the meaning of predictors is the same as in the DTA model. However, migration is parameterized as a backwards-in-time migration rate matrix (*m*) and the effective population size 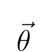 is modelled separately for each sub-population. The first term is the likelihood of sequences given genealogy and substitution model, which is computed using the pruning algorithm (Felsenstein, 1981). The second term is the probability density of the genealogy and migration history of lineages under the structured coalescent assumption given migration rate matrix and effective population sizes. The third term represents the prior distribution of the model parameters.

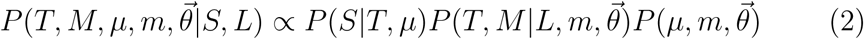

The structured coalescent likelihood can be computed analytically only when conditioned on a migration history (*M*). Thus standard approaches have required augmenting the tree with a random-dimensional migration history (M) which has restricted the application of this model to datasets with a small number of demes/locations (Vaughan et al., 2014). However, when the details of the migration history are not of particular interest, the MASCOT model (Müller et al., 2017) can be used to approximate the integrated likelihood (i.e. formally integrating over every possible migration history for each tree in the MCMC chain). This approximation is closely related to the exact structured coalescent, but still allows us to analyse dataset with many different sub-populations. Since we seek to investigate the spread of H9N2 between 12 different countries and regions, we used MASCOT in our analyses.

### Migration rate GLMs

DTA and MASCOT have both been extended such that constant migration rates and time-series migration rates can be described using GLMs (Lemey et al., 2014; Bielejec et al., 2014; Müller et al., 2017). This allows us to infer the contribution of explanatory factors to migration rates between different sub-populations, and through time. Both the prior on inclusion probability of predictor and the effect of including isolate sample size as a predictor, were separately examined in our GLM models. We used a strict and weak prior probability on the inclusion of each predictor to reflect 50% and 5% prior probability on no predictors being included in the GLM respectively, and the prior probability on each predictor’s inclusion is equal. Four variants were considered in GLM under DTA model to investigate contribution of drivers to the constant migration rates among these unstructured subpopulations; both time-dependent and time-independent predictors were considered to model viral migration rates between different sub-populations in eight GLMs using MASCOT.

Migration rates between locations are defined as log-linear combinations of coefficients, indicators and predictors. Eq 3 and 4 describe the time-independent and time-dependent parameterizations of the migration rates respectively. Here, *m_ij_* represents the migration rates between location *i* and *j*; 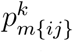 represents the kth predictor between location *i* and *j;* 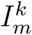 represents the indicator and 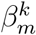 represents the coefficient of the kth predictor. The indicator and coefficient describe if and to what degree each predictor contributes to explain the migration rates. Indicators are estimated by a Bayesian stochastic search variable selection (BSSVS) algorithm to describe the posterior inclusion probability for each predictor and use the priors on the number of active predictors to reduce over-fitting (Chipman et al., 2001). Further, *m_ij_*(*t*), 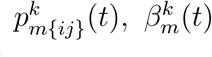 and 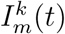 represent the time-dependent version of corresponding parameters in this model. Because more than 10 predictors were chosen to model the migration rates, we removed the error terms in the GLM model for fear of overparameterization.

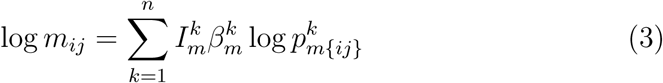

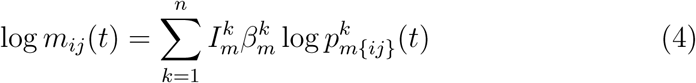

In contrast to DTA, the structured coalescent is parameterized by backwards in time migration rates, 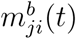. These backwards in time migration rates we get by scaling the forward in time migration rates, 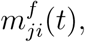, with effective population sizes of the source and sink population, such that the backwards in time rates are:

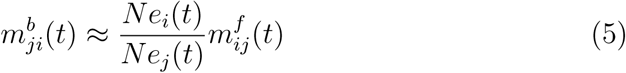

### Population dynamic GLMs

Effective population sizes within different sub-populations were modelled by both time-independent (Eq 6) and time-dependent (Eq 7) GLM models in MASCOT. Here, *Ne_i_* represents the viral effective population size of location 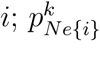 represents the *k*th (time-independent) predictor at location 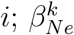 and 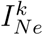 represent the coefficient and inclusion probability of the *k*th predictor respectively. *α_i_* represents the extra part of viral effective population size that could not be explained by the predictors in region *i*. Further, *N_ei_*(*t*), 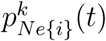,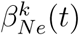 and 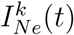 represent the time-dependent version of corresponding parameters from 1986 to 2013 in the model.

Eq 8 describes a specific instance of the model in Eq 7. The first part of Eq 8 describes the relationship between viral effective population size and poultry production in each region jointly. Here, the number of predictors *n* is equal to the number of locations in the analysis. 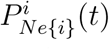 represents the poultry production in region *i* from 1986 to 2013. In order to model the time before 1986 as a structured coalescent process with constant rates, we introduce predictors that are 1 for any of the locations before 1986 and 0 otherwise. One predictor therefore only predicts the *Ne* after 1986 and for only one location. This we do in order to avoid events that happened more than 28 years ago for which we do not have predictor information to bias our inference.

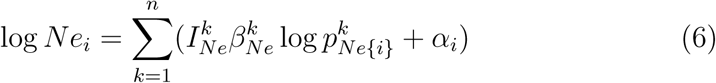

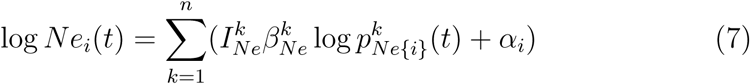

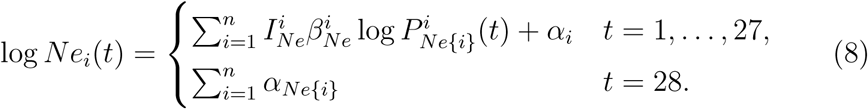

### Parameter inference

Bayesian analyses of H9N2 AIVs using the DTA and DTA GLM models were conducted using BEAST v1.10.0 (Suchard et al., 2018). The MASCOT GLM analyses on the same data were conducted using BEAST v2.5.0 (Bouckaert et al., 2014). An HKY nucleotide substitution model with gamma site heterogeneity using 4 rate categories and a strict molecular clock model were employed to model sequence evolution in all analyses. The discrete phylogeographic analysis using DTA with a symmetric trait substitution model and Bayesian skyline tree prior was performed in 5 parallel runs. The convergence and mixing of MCMC chains in these runs were diagnosed by the RWTY package in R v3.4.3 (Warren et al., 2017). DTA and MASCOT GLM models were used to estimate the contribution of potential predictors to the migration rates between each pair of locations. Further, the MASCOT analyses included population dynamic GLMs to identify the underlying factors driving virus population diversity in each sub-population. Although MASCOT improved the computational efficiency of the structured coalescent, it is still a more computationally demanding analyses than the DTA model. Therefore, in models using MASCOT, we fixed the phylogenetic tree to the maximum clade credibility (MCC) tree obtained from the DTA analysis. We performed at least 5 runs with 10-100 million iterations (based on the convergence time of different analyses) of different analyses to estimate the phylogenetic tree with location information and GLM parameters. We used Tracer v1.7 (Rambaut et al., 2018) to remove an appropriate burn-in (10%-20% of samples in most cases) to achieve an adequate effective sample size (ESS, at least 100). The time scaled phylogenetic tree with the maximum probable location in each lineage was annotated and visualized in FigTree v1.4.3 (http://tree.bio.ed.ac.uk/software/figtree/) and the ggtree package in R v3.4.3 (Yu et al., 2017).

### Data availability

The original H9N2 HA nucleotide sequences and all predictors considered in this study as well as the BEAST xml files are available at https://github.com/judyssister/avianInfluenzaH9N2.git. The log files on estimated parameters and MCC tree files generated from these analyses are also available in the same GitHub repository.

## Results

### H9N2 was endemic in domestic poultry in Asian countries

H9N2 viruses spread to at least 22 countries in Asia between 1976 and 2014 (Fig 1a). These 22 countries are mostly located in tropic and temperate zones with lower latitude, where the climatic conditions are suitable for poultry rearing. H9N2 viruses have been isolated from a wide variety of different hosts, including the major poultry species: chickens and ducks. Compared to the number of isolates from wild birds, H9N2 viruses were predominantly isolated from domestic poultry populations in Asia, especially from chicken. Since 1996, H9N2 viruses have been isolated in more Asian countries and then persisted in birds in some of these countries for several years (Fig 1b). The number of isolates shows an increasing trend and most were sampled from mainland China since the late 1990s, which is likely in part driven by a larger surveillance effort. The estimated effective population size of the virus however also substantially increased since 1996 (Fig 1c).

**Figure 1:**
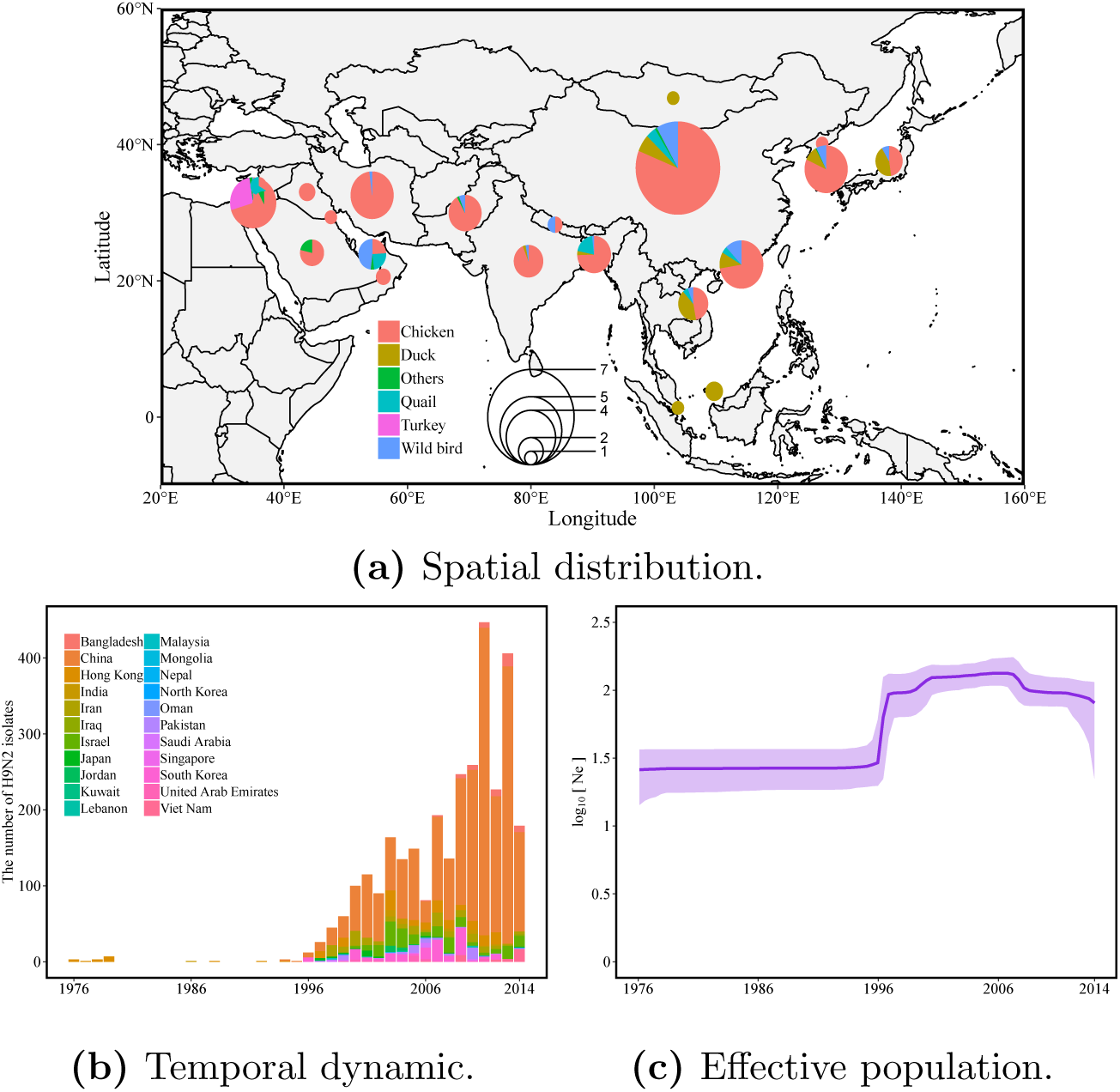
Spatio-temporal distribution of avian-origin H9N2 viruses in Asia between 1976 and 2014. (a) Spatial distribution where the size of circle represents the number of virus isolates obtained in the corresponding countries/locations. The value of circle indicates double square roots of virus isolate quantity in each location. Colors in the circle represent different fraction of virus host origins. Most of the isolates originate from chickens and from China. (b) Number of H9N2 HA isolates that are deposited in NCBI through time. The bars represent the annual number of H9N2 isolates sampled in each region. (c) Inferred effective population sizes of H9N2 in Asia by using Bayesian skyline plot. The effective population size of H9N2 viruses increased in 1996 when more isolates began to be sampled in larger number countries representing multiple outbreaks.

### Viral phylogeny and dynamics of spread

The evolutionary relationships and the migration history of lineages reconstructed using the two phylogeographic methods are shown in Fig 2. As mentioned previously, the genealogies can be grouped into three lineages: G9-like, G1-like and Korea-like lineage (Guo et al., 2000). These three lineages established in 1990s and continued to be isolated to date. Most isolates used in this study were found to be G9- or G1-like. Isolates from countries close by to each other were often genetically related (Fig 2). G9-like H9N2 viruses were mainly isolated in mainland China, Hong Kong, Japan, and Vietnam. G1-like viruses were mainly isolated in the Middle East and the Indian subcontinent. Korea-like viruses were predominantly isolated in South Korea, Japan, and Hong Kong.

Hong Kong was inferred as the most likely source of H9N2 viruses circulating in Asia by both DTA and MASCOT (Fig 2). DTA inferred H9N2 to have spread from Hong Kong to East Asia in the 1980s. After, one part of the viral population continued to spread in countries in the East and Southeast Asia; Another part of the population was transmitted to several countries in the West and South Asia. MASCOT inferred that H9N2 viruses were directly transmitted into countries in the West and South Asia from Hong Kong, and via multiple introductions from Hong Kong to East Asia.

Viral migration rates among Asian countries and regions were inferred by using DTA with symmetric migration rates (Fig 3). The highest migration rates with the strongest support were inferred between the United Arab Emirates and Saudi Arabia, between Hong Kong and Vietnam, between Iran and Pakistan, between Hong Kong and mainland China, between the United Arab Emirates and Pakistan, and between mainland China and Japan. All these pairs are locations within close geographic proximity. The migration rates among these six pairs are 0.92 or higher, which means ∼ 1 migration event occurred between these locations per lineage per year.

**Figure 2:**
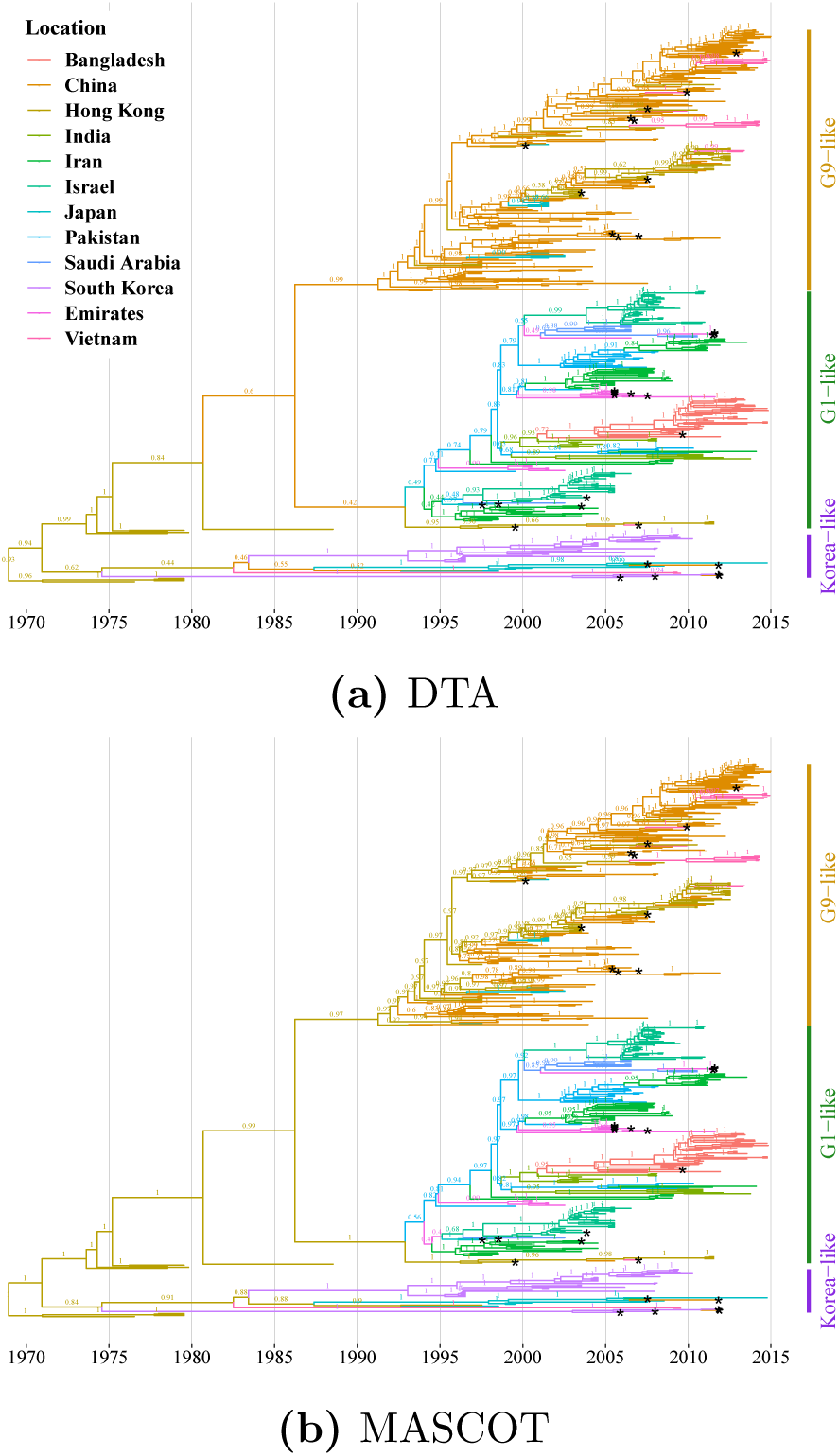
Time scaled phylogenetic trees of H9N2 virus in Asia. (a) estimated using DTA model and (b) using MASCOT model. The colour of tree branches indicates location (see legend) with the maximum probability. A colour change on a branch indicates a virus migration event. Numbers on branches represent posterior probability of the displayed location. A black asterisk represents a virus sequence isolated from the wild bird. Both methods place the origin of H9N2 in Hong Kong, from where it spread to East Asia. DTA and MASCOT differ in the details on how it spread to West and South Asia. Bars on the right indicate three established lineages based on the phylogenetic relationship between the virus and the representative strains in Asia. The phylogenetic cluster of isolates from domestic poultry in nearby 18 regions indicates their roles in virus spread among neighbouring locations; whereas the dispersal distribution of isolates from wild birds on the phylogeny questions their roles in virus spread across countries and genetic groups.

**Figure 3:**
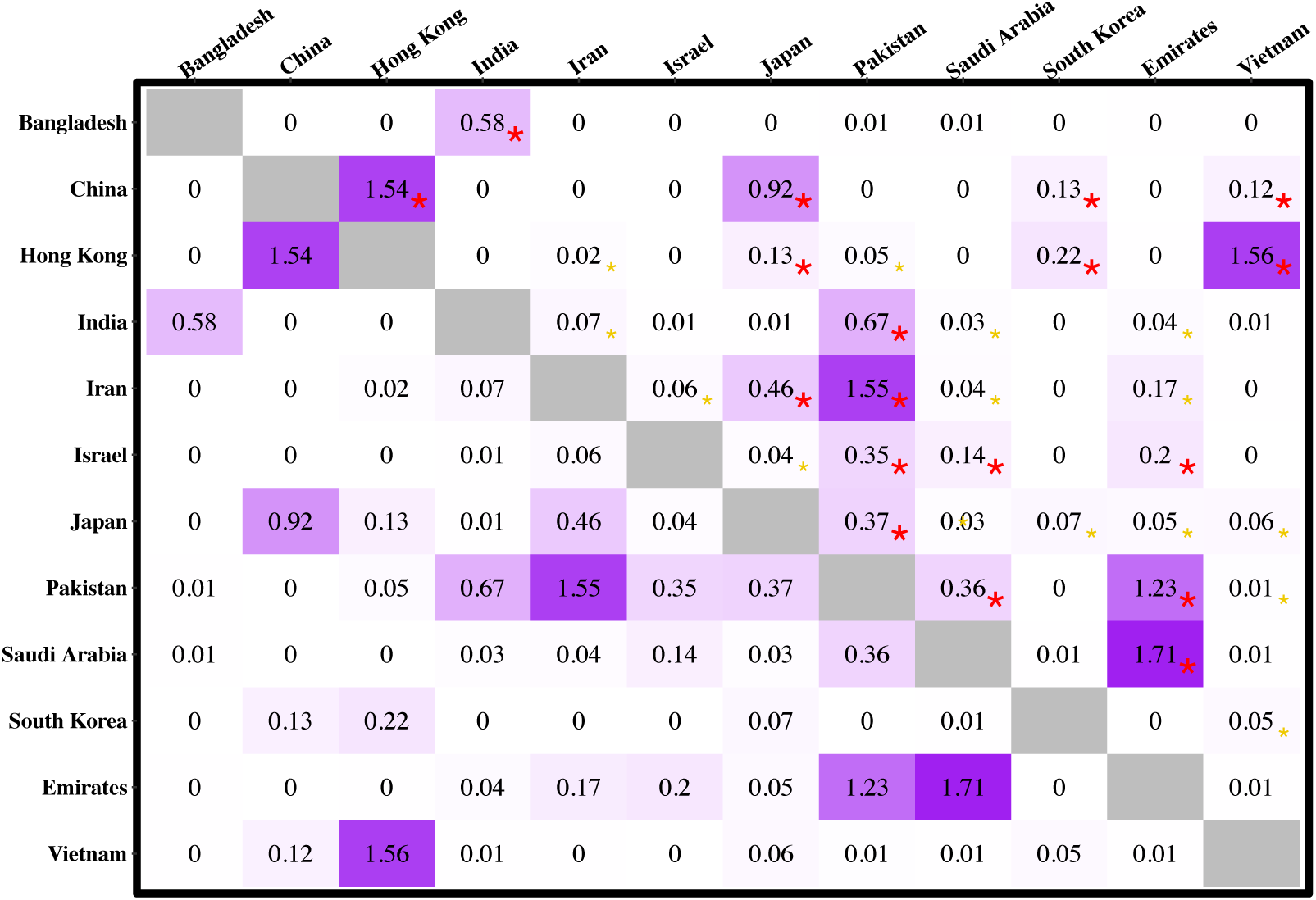
Symmetric migration rate matrix of H9N2 viruses between each pair of locations in Asia. The migration rate matrix was estimated using DTA. The matrix describes the virus spread between each pair of locations without migration direction. Units are the number of migration events per lineage per year. Bayes factors on migration rate over 3 and 20 are labeled by a yellow and a red asterisk in the upper triangular matrix respectively. The largest and most well-supported rates are between neighbouring locations, suggesting the underlying factors related to geographic proximity could contribute to virus spread.

### The mechanisms of virus spatial spread

We next investigated the factors driving the spread of H9N2 viruses across several locations in Asia by using a GLM to model the relationship between potential predictors and viral migration rates (Fig 4). A total of 12 different GLMs were used S2 and S3 Tables. We used both DTA and MASCOT and included time-dependent and time-independent predictors; strict or weak prior probabilities on the inclusion probability of each predictor. Further, we ran analyses with and without considering the number of isolates as a distinct predictor. Bayes factors (BFs) on inclusion probability of each predictor were calculated to explain how much the data informed the inclusion of a predictor (Suchard et al., 2001). BF is calculated as a ratio of the posterior odds for a predictor inclusion to the corresponding prior odds. A BF over 3 is typically considered suggestive and a BF over 20 is typically used as strong supporting a predictor to be included into the GLM model (Kass and Raftery, 1995). Geographic distance was identified as a strong supported driver in all migration rate GLMs investigated and it consistently made a negative contribution to the virus spread (S3 Table), meaning that migration is inferred to be stronger between closer countries. Predictors related to rainfall seasonality in destination location, poultry trade and if two locations sharing border on continent were inferred in more than half of these 12 GLMs investigated to be strongly supported. Rainfall seasonality in destination showed a negative relationship with the virus migration rates. This suggests that the virus could more easily spread to countries and in years with less rainfall variation throughout the year. Higher poultry trade and sharing a border were both positively associated with virus spread. These results were consistent and robust when we repeated the analysis whether considering virus sample size as a distinct predictor or giving a strict or weak prior to indicator inclusion.

**Figure 4:**
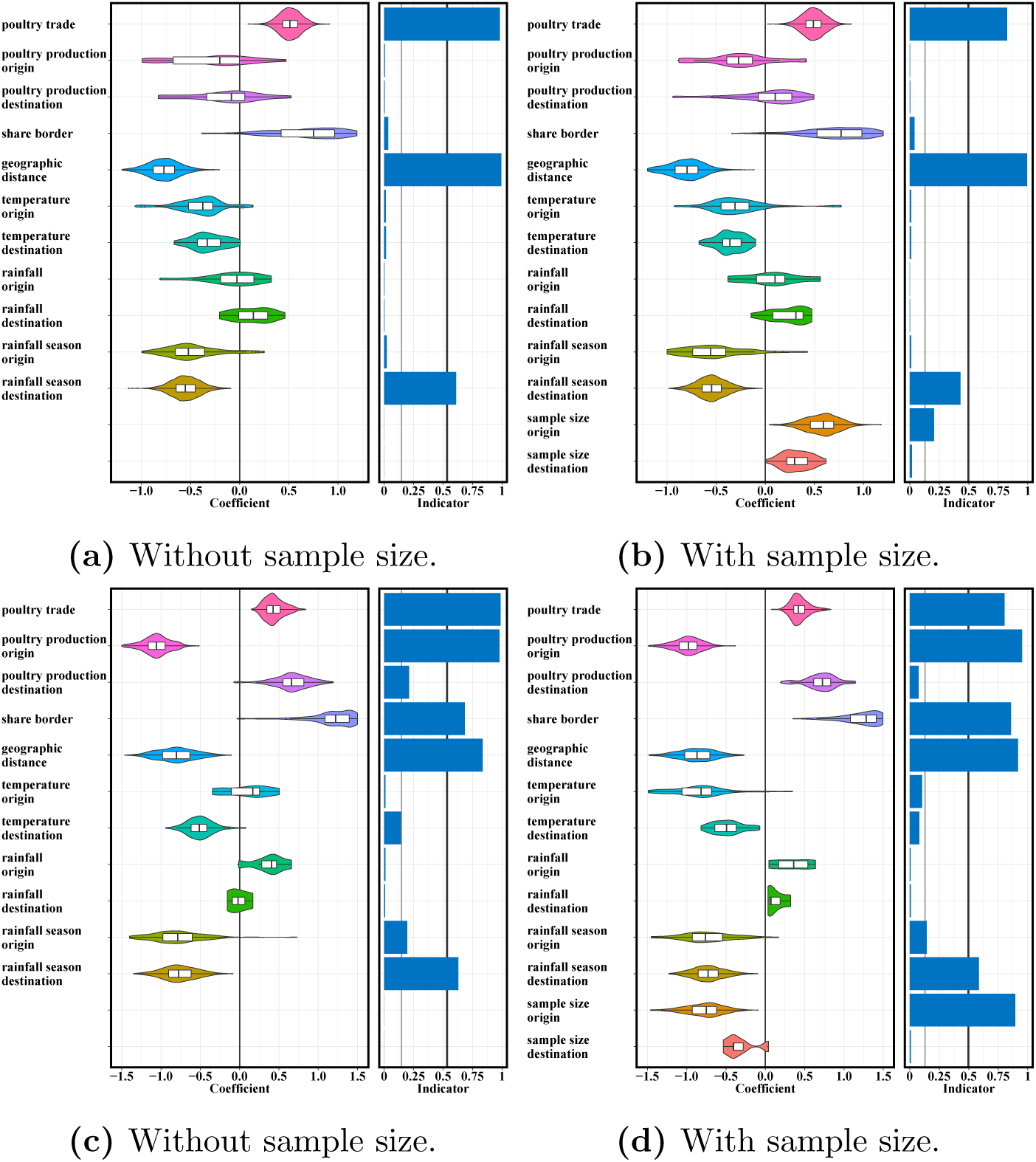
Predictors of H9N2 migration rates between 12 countries/locations in Asia. The estimated coefficients and inclusion probabilities for potential predictors of migration rates in the DTA model: (a) without and (b) with isolate sample sizes included as potential predictors; in the time-dependent MASCOT GLMs: (c) without and (d) with isolate sample sizes considered as a predictor. The 50% prior mass was specified on no predictors being included. Coefficients represent the contribution of each predictor to the migration rates of H9N2 AIVs when the corresponding predictor was included into the model. Inclusion probabilities are calculated by proportion of the posterior samples in which each predictor was included in the model. Bayes factor support v2a1lues of 3 and 20 are represented by a thin and thick vertical line respectively in the inclusion probabilities plot. Geographic distance, poultry trade and rainfall seasonality in destination are the most strongly supported factors to virus spread in Asia under cross-validation in these models. Sample size at origin has an effect, but it doesn’t change the support of other predictors.

### Predictors of viral effective population sizes through time

In MASCOT, we also jointly inferred predictors of effective population sizes of H9N2 virus in the different locations and their changes through time. The effective population sizes in the different locations were modelled by multiple time-independent and time-dependent predictors in GLMs. Poultry production was selected as a suggestive predictor for the effective population size of H9N2 virus in GLM model with time-dependent predictors (Fig 5). This implies that the higher the poultry production in a country is, the larger the genetic diversity of H9N2 virus in that country. Furthermore, virus sample size was also considered as a positively supportive predictor and the inclusion of virus sample size into GLM model lowered the contribution of poultry production to virus population diversity. This suggests that the number of viral samples through time was roughly proportional to the effective population sizes. No potential predictors of effective population sizes were supported when we assumed that the effective population sizes are constant through time. Additionally, we applied a GLM that used time-dependent poultry production data in 12 Asian countries or regions to model the virus population dynamics in each subpopulation jointly. Poultry production in mainland China was strongly supported as a positive predictor of the local viral effective population size. We also found suggestive support for poultry production in Bangladesh, but not in any other location.

**Figure 5:**
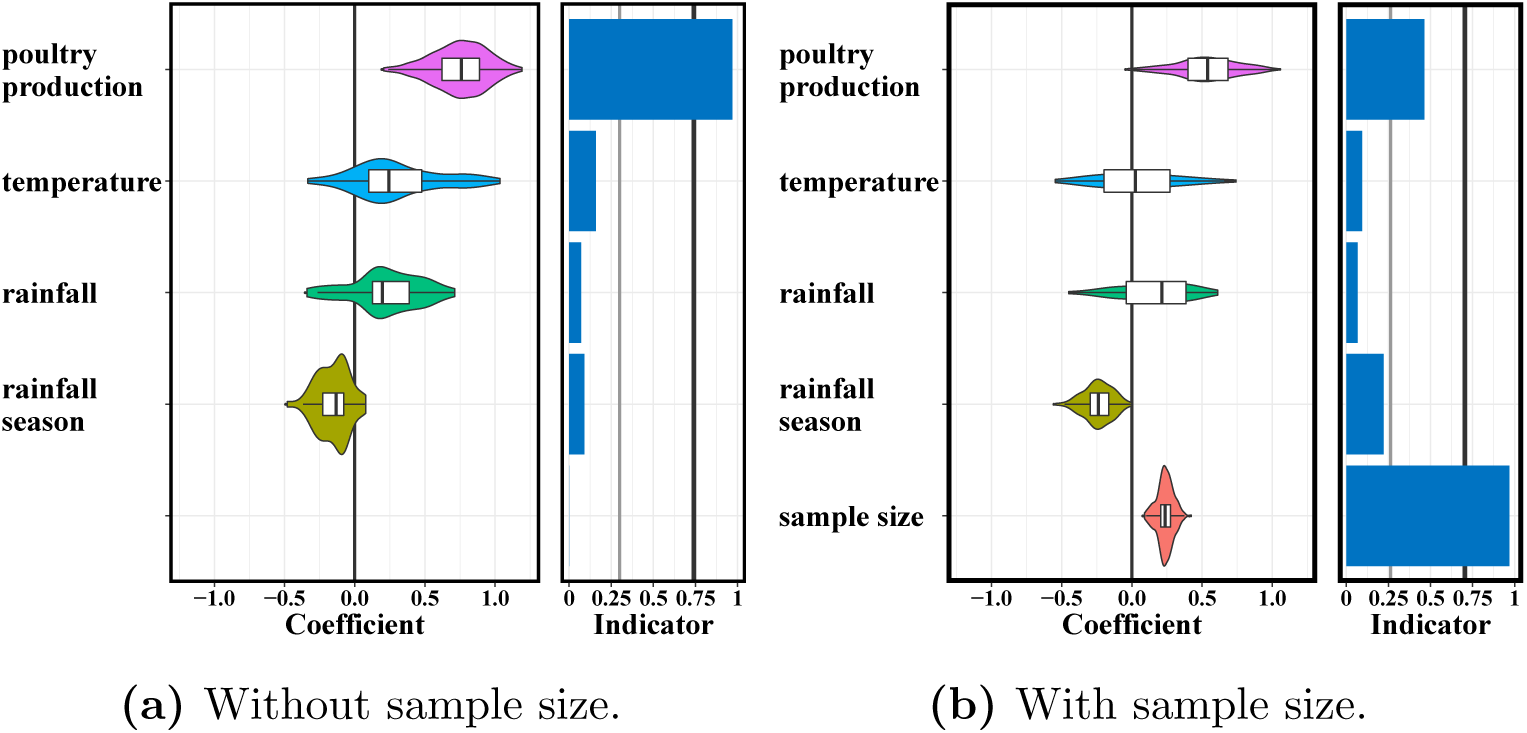
Predictors of time-dependent H9N2 population dynamics within 12 countries/locations in Asia. Coefficients and indicators as in Fig 4 when estimating the effective population size of H9N2 AIVs (a) without and (b) with considering the effect of virus sample size included as a distinct predictor. The 50% prior mass was specified on no predictors being included. Bayes factor support values of 3 and 20 are represented by a thin and thick vertical line respectively in the indicator plot. Poultry production positively contributes to virus population size. When the number of samples through time in each location is also used as a predictor, the effect of poultry production is much less pronounced; we questions that our sampling may have been approximately proportional to effective population sizes.

## Discussion

In Asia, H9N2 viruses have spread into multiple countries which did not previously have documented viral isolation in the 1990s and persist in domestic poultry in some of these countries (Guan et al., 1999; Cameron et al., 2000; Nili and Asasi, 2003). To reduce the cost and maximize the profit, poultry farms are often built in close proximity to one another and rearing facilities tend to be overcrowded (Mehrabadi et al., 2018). Viruses can therefore spread easily and outbreaks in poultry pose great economic threats to some of these countries. Further, most of these countries are developing countries with poor bio-security. Low sanitary standards and high density of poultry in farms and markets can additionally facilitate the transmission of viruses (Gao, 2014). Multiple influenza subtypes simultaneously circulate in birds in these countries (Shanmuganatham et al., 2014; Thuy et al., 2016), which increases the probability of reassortment of influenza segments.

In this work, we investigated the evolutionary dynamics and the spread of avian influenza H9N2 in Asia and attempted to uncover factors that potentially predict this spread by using a GLM in two phylgeographic frameworks, DTA and MASCOT (Lemey et al., 2014; Müller et al., 2018). Alongside estimating the factors driving migration rates, we also jointly investigated potential drivers of virus effective population sizes in MASCOT. To do so, we used H9N2 viral HA sequences isolated from avian hosts and 12 locations in Asia between 1976 and 2014. We used different predictor data to inform the viral migration rates between 12 countries/locations. These predictors however ignore other potential drivers of migration, such as wild bird migration, and different sanitation levels among countries. Typically, predictors adopted to predict the virus spread and diversity were scale-dependent. In the future, more exact and more high-resolution predictors could be included to test more detailed hypotheses and model influenza movements in a smaller and confined geographical region (Lemey et al., 2014).

Since rate estimates of DTA are likely to be sensitive to the number of sequences sampled in each location (De Maio et al., 2015), we performed analyses with and without considering the viral isolate sample size in each country or region as a distinctive predictor. The results were mostly robust and consistent whether this predictor was included or not in DTA. But the inclusion of isolate sample size in the model did reduce the support of predictor about rainfall seasonality in the destination location, which is negatively related to virus migration rates. Due to computational reasons, we used a fixed phylogenetic tree in MASCOT, meaning that we ignored phylogenetic uncertainty in that analysis.

We found geographic proximity between locations to be a strong driver of H9N2 migration rates in all GLM models investigated. Additionally, we found that whether two locations are neighbouring each other to be a strong predictor of migration. The contribution of geographic proximity to viral spread was intuitively recorded in the close evolutionary relationship among viruses sampled from nearby countries. Further, a consistent role for poultry trade was inferred in both a GLM with time-independent predictors in DTA and a GLM with time-dependent predictors in MASCOT. But this variable did not have strong support in the GLM model with time-independent predictors in MASCOT, showing that inferences are sensitive to model assumptions. This suggests that poultry trade is partly responsible for the spread of avian influenza H9N2 viruses. Infected poultry, especially chicken, without strong clinical symptoms can easily be missed during process of transportation. H9N2 viruses can therefore spread into native poultry. Increased surveillance of imported poultry and their products could decrease the spread of H9N2 across locations (MASE et al., 2007). Illegal poultry trade across borders is another potential factor contributing to the spread of H9N2 (Thuy et al., 2016). However, even when controlling for poultry trade volumes and other potential predictors, we still found geographic proximity to be a key driver to migration rates. This may point to some factors directly linked to geographical distance to contribute to the viral spread of H9N2 across countries. Contact between domestic and wild birds is inevitable in the transitional and intensive livestock and two-way virus transmission has been documented between them (Bahl et al., 2016; Gu et al., 2017). Wild birds could therefore spread H9N2, as they can easily cross borders and then transmit the viruses. The dispersal distribution of H9N2 isolates from wild birds on the phylogenetic tree supports the possibility of their movement facilitating virus spread across countries and across genetic groups (Fig 2). Future studies will however have to investigate if wild bird migration is really associated with the spread of H9N2 viruses. Further, the region with less monthly variance in rainfall volumes could provided a stable forging and habitat areas for birds and attracted the birds carrying viruses. Active surveillance of migratory birds could therefore help to monitor the dispersal of H9N2 virus.

To improve our understanding of what potentially drives genetic diversity of H9N2, we also used a GLM approach to inform effective population sizes of H9N2 virus in each location and through time by using MASCOT (Müller et al., 2018). Time-dependent poultry production was identified as a positive driver to virus divergence within each sub-population. When including the number of samples through time, the support for poultry production as a effective population size predictor however decreased. This can be caused by a proportional relationship between the annual number of viral samples and the viral effective population size in a location changed over time. The viral genetic diversity was at least partly driven by poultry production in mainland China and Bangladesh. Approximately 78% of H9N2 samples were isolated from China based on the HA gene sequences recorded in NCBI (Li et al., 2017). The positive correlation between an increasing poultry production and an increasing effective population size of the virus suggests that virus control measures in local poultry may not currently be sufficient in these countries.

Surveillance of high pathogenic H5N1 AIVs in ducks has been actively carried out in several Asian countries (Nguyen et al., 2009). Samples from chickens, wild bird, and the environment could also be collected to investigate the prevalence H9N2 and other subtypes. If there are H9N2 cases in humans, having such samples readily available can help to track possible origins of these cases. Additionally, making surveillance of both high pathogenic and low pathogenic AIVs in poultry and humans routine can potentially help to improve our understanding of how these viruses jump into humans. Overall, the integration of temporal predictors into phylodynamics provides a powerful tool to test how disease spread within and between populations.

## Acknowledgments

We would like to thank the China Scholarship Council (CSC) for the financial support to Jing Yang to visit the Centre of Computational Evolution in University of Auckland, New Zealand to carry out this work. We also would like to thank the New Zealand eScience Infrastructure (NeSI) for their supply of high-performance computing facilities. JY is funded by the National Key Research and Development Program of China (NO.2016YFA0600104). NFM is funded by the Swiss National Science foundation (SNF; grant number CR32I3 166258). AJD would like to acknowledge support from a Royal Society of New Zealand Marsden award (#UOA1611; 16-UOA-277).

